# Systemic characterization of pppGpp, ppGpp and pGpp targets in *Bacillus* reveals NahA converts (p)ppGpp to pGpp to regulate alarmone composition and signaling

**DOI:** 10.1101/2020.03.23.003749

**Authors:** Jin Yang, Brent W. Anderson, Asan Turdiev, Husan Turdiev, David M. Stevenson, Daniel Amador-Noguez, Vincent T. Lee, Jue D. Wang

## Abstract

The alarmones pppGpp and ppGpp (collectively (p)ppGpp) protect bacterial cells from nutritional and other stresses. Here we demonstrate the physiological presence of pGpp as a third closely related alarmone in bacterial cells and also characterize and compare the proteomic targets of pGpp, ppGpp and pppGpp in Gram-positive *Bacillus* species. We revealed two regulatory pathways for ppGpp and pppGpp that are highly conserved across bacterial species: inhibition of purine nucleotide biosynthesis and control of ribosome assembly/activity through GTPases. Strikingly, pGpp potently regulates the purine biosynthesis pathway but does not interact with the GTPases. Importantly, we identified a key enzyme NahA that efficiently produces pGpp by hydrolyzing (p)ppGpp, thus tuning alarmone composition to uncouple the regulatory modules of the alarmones. Correspondingly, a *nahA* mutant displays significantly reduced pGpp levels and elevated (p)ppGpp levels, slower growth recovery from nutrient downshift, and loss of competitive fitness. These cellular consequences for regulating alarmone composition strongly implicate an expanded repertoire of alarmones in a new strategy of stress response in *Bacillus* and its relatives.

## Introduction

Organisms from bacteria to humans rely on timely and appropriate responses to survive various environmental challenges. The stress signaling nucleotides guanosine tetraphosphate (ppGpp) and guanosine pentaphosphate (pppGpp) are conserved across bacterial species. When induced upon starvation and other stresses, they mediate multiple regulations and pathogenesis by dramatically remodeling the transcriptome, proteome and metabolome of bacteria in a rapid and consistent manner^1–3^. (p)ppGpp interacts with diverse targets including RNA polymerases in *Escherichia coli* ^4–8^, replication enzyme primase in *Bacillus subtilis*^9–11^, purine nucleotide biosynthesis enzymes^12–15^, and GTPases involved in ribosome assembly^16–19^. Identification of (p)ppGpp binding targets on a proteome-wide scale is one way to unravel a more extensive regulatory network^15,18,20^. However, because binding targets differ between different species and most interactomes have not been characterized, the conserved and diversifying features of these interactomes remain incompletely understood.

Another understudied aspect of (p)ppGpp regulation is whether ppGpp and pppGpp, while commonly referred to and characterized as a single species, targets the same or different cellular pathways^21^. In addition, there is evidence for potential existence of a third alarmone, guanosine-5’-monophosphate-3’-diphosphate (pGpp), since several small alarmone synthetases can synthesize pGpp *in vitro*^22,23^. However, the clear demonstration of pGpp in bacterial cells has been challenging. More importantly, the regulation specificities and physiological importance of having multiple closely-related alarmones in bacteria have not been systematically investigated.

Here we demonstrate pGpp as a third alarmone in Gram-positive bacteria by establishing its presence in cells, systematically identifying its interacting targets, and revealing a key enzyme for pGpp production through hydrolyzing (p)ppGpp. We also compare the targets of pGpp, ppGpp and pppGpp through proteomic screens in *Bacillus anthracis*. We found that both pppGpp and ppGpp regulate two major cellular pathways: purine synthesis and ribosome biogenesis. In contrast, pGpp strongly regulates purine synthesis targets but does not regulate ribosome biogenesis targets, indicating a separation of regulatory function for these alarmones. In *B. subtilis* and *B. anthracis*, pGpp is efficiently produced from pppGpp and ppGpp by the NuDiX (<u>Nu</u>cleoside <u>Di</u>phosphate linked to any moiety “<u>X</u>”) hydrolase NahA (<u>N</u>uDiX <u>a</u>larmone <u>h</u>ydrolase A), both *in vitro* and *in vivo*. A Δ*nahA* mutant has significantly stronger accumulation of pppGpp and decreased accumulation of pGpp, as well as slower recovery from stationary phase and reduced competitive fitness against wild type cells. Our work suggests a mechanism for the conversion and fine tuning of alarmone regulation and the physiological production of the alarmone pGpp.

## Results

### Proteome-wide screen for binding targets of pppGpp and ppGpp from Bacillus anthracis

To systematically characterize the binding targets of (p)ppGpp and identify novel (p)ppGpp binding proteins in *Bacillus* species, we screened an open reading frame (ORF) library of 5341 ORFs from the pathogen *Bacillus anthracis* (**Figure 1a**). Using Gateway cloning, we placed each ORF into two expression constructs, one expressing the ORF with an N-terminal histidine (His) tag and the other with an N-terminal histidine maltose binding protein (HisMBP) tag.

**Figure 1.**
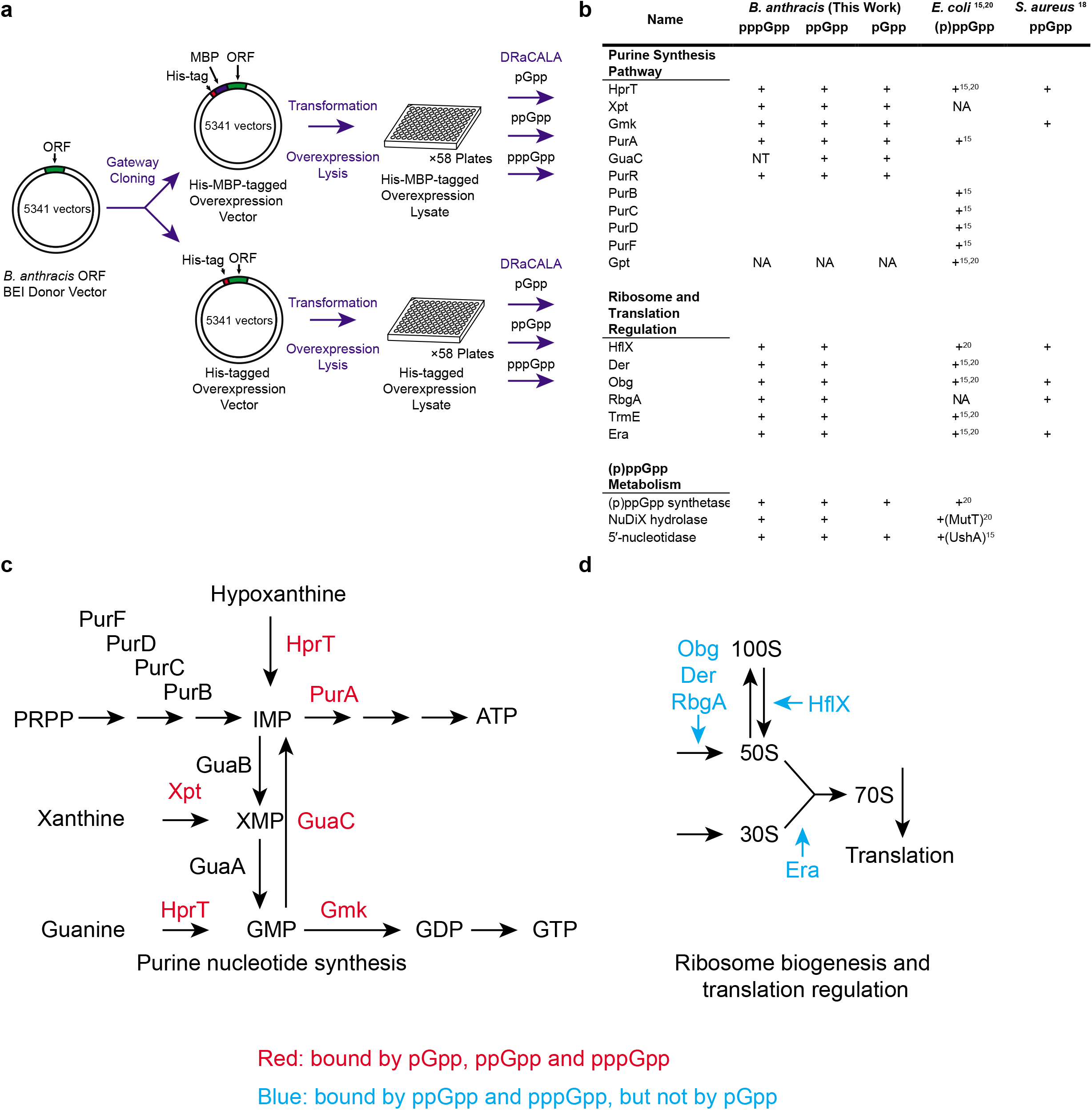
Proteome-wide DRaCALA screen identifies both conserved categories of binding targets and novel targets. **a**) The *Bacillus anthracis* ORF donor vector library was recombined by Gateway cloning into overexpression vectors to generate ORFs with an N-terminal His-tag or His-MBP tag. The plasmids were transformed into *E. coli* for overexpression of recombinant proteins. Lysates of each ORF overexpressed in *E. coli* were assayed for binding to pppGpp, ppGpp and pGpp using DRaCALA. **b**) List of identified (p)ppGpp binding targets in *E. coli, S. aureus*, and *B. anthracis* and pGpp binding targets in *B. anthracis*. **c-d**) Schematics of pathways differentially regulated by pGpp and (p)ppGpp: **c**) Enzymes in purine nucleotide synthesis, including HprT, Xpt, Gmk and GuaC, bound both pGpp and (p)ppGpp; **d**) GTPases, involved in ribosome biogenesis and translational control (Obg, HflX, Der, RbgA and Era), bound (p)ppGpp, but not pGpp.

We first charaterized the binding targets of ppGpp using the *B. anthracis* library. To this end, each ORF in the His-tagged and HisMBP-tagged library was overexpressed and binding [5’-α-^32^P]-ppGpp was assayed using differential radial capillary action of ligand assay (DRaCALA)^24^ (**Figure 1a**). The fraction of ligand bound to protein in each lysate was normalized as a Z-score of each plate to reduce the influence of plate-to-plate variation (Table S1). We found that the strongest ppGpp-binding targets in *B. anthracis* can be categorized to three groups: 1) purine nucleotide synthesis proteins (Hpt-1, Xpt, Gmk, GuaC, PurA, PurR); 2) ribosome and translation regulatory GTPases (HflX, Der, Obg, RbgA, TrmE, Era); and 3) nucleotide hydrolytic enzymes, including NuDiX hydrolases and nucleotidases (**Figure 1b**). We compared these targets to those obtained from previous screens for ppGpp targets in *E. coli* and for an unseparated mix of pppGpp and ppGpp in *S. aureus*^18^. Comparison of our results with these previous screens yielded conserved themes (**Figure 1b**). Among the most conserved themes are the purine nucleotide synthesis proteins (**Figure 1c**) and ribosome and translation regulation GTPases (**Figure 1d**).

Next, we performed a separate screen to characterize the binding of the *B. anthracis* proteome to pppGpp (**Figure 1a**). pppGpp is the predominant alarmone induced upon amino acid starvation in *Bacillus* species, rising to a higher level than ppGpp. However, despite potential differences in specificity between pppGpp and ppGpp, the pppGpp interactome has not been systematically characterized in bacteria. We found that pppGpp shares almost identical targets with ppGpp, with similar or reduced binding efficacy for most of its targets compared to ppGpp (**Table S1**). By sharing targets with ppGpp, pppGpp also comprehensively regulates purine synthesis and ribosome assembly. We also find that several proteins bind to pppGpp but not ppGpp, including the small alarmone synthetase YjbM (SAS1). This is expected for YjbM, since it is allosterically activated by pppGpp, but not ppGpp^25^.

### NahA, a NuDiX hydrolase among the (p)ppGpp interactome in *Bacillus*, hydrolyzes (p)ppGpp to produce pGpp *in vitro*

The putative NuDiX hydrolase, BA5385, was identified as a novel binding target of (p)ppGpp. Protein sequence alignment showed that BA5385 has homologs in different *Bacillus* species with extensive homology and a highly conserved NuDiX box (**Figure S1**). We cloned its homolog, YvcI, from the related species *Bacillus subtilis* and showed that overexpressed *B. subtilis* YvcI in cell lysate also binds ppGpp and pppGpp (**Figure 2a**). The binding is highly specific, as non-radiolabeled ppGpp effectively competes with radiolabeled (p)ppGpp binding, whereas non-radiolabeled GTP failed to compete. EDTA eradicated (p)ppGpp binding to His-MBP-YvcI cell lysate, which implies that the divalent cations present in the reaction (Mg^2+^) is essential for (p)ppGpp binding to YvcI (**Figure 2a**).

**Figure 2.**
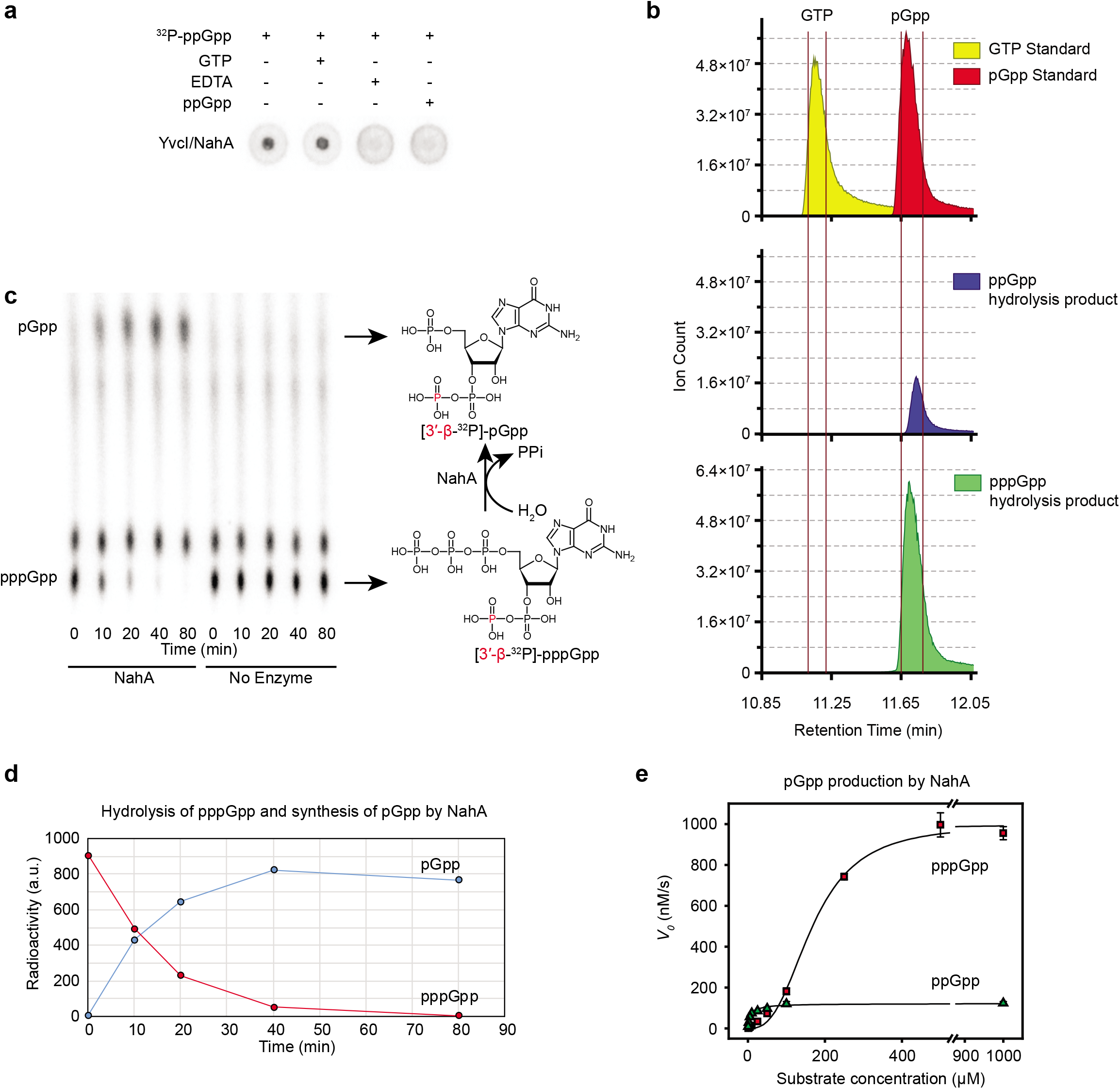
NahA (YvcI) produces pGpp via (p)ppGpp hydrolysis. **a**) DRaCALA of [5’-α-^32^P]-ppGpp binding to *B. subtilis* HisMBP labeled -YvcI (NahA) overexpressed in *E. coli* cell lysate. Unlablled GTP (100 uM), unlabeled ppGpp (100 μM) and EDTA (10 mM) were added as indicated. **b**) Ion count vs. retention time curves from LC-MS of GTP and pGpp standards and NahA-catalyzed hydrolysis products from ppGpp and pppGpp. **c**) TLC analysis of NahA activity over time with [3’-β-^32^P]-pppGpp. Expected NahA-catalyzed conversion of [3’-β-^32^P]-pppGpp to [3’-β-^32^P]-pGpp is shown on the right. Radiolabeled [3’-β-^32^P]-phosphorus atom is highlighted in red. **d**) Quantitation of pppGpp and pGpp in the hydrolysis of pppGpp in (c). **e**) Initial velocity vs. pppGpp or ppGpp concentration curves for NahA synthesis of pGpp. Data were obtained by kinetic assay using radiolabels (see Methods). Curves represent the best fit of the data from three independent experiments. Error bars represent standard error of the mean.

We noticed that YvcI overexpression cell lysate showed strong and specific binding to (p)ppGpp, the purified protein does not appear to bind (p)ppGpp in DRaCALA (**Figure S2**). This suggests that either YvcI requires a co-factor present in the lysate to bind to (p)ppGpp, or YvcI may rapidly hydrolyze the (p)ppGpp and release the product. Therefore, we incubated purified YvcI with [5’-α-^32^P]-(p)ppGpp and ran the reaction product using TLC (**Figure S3**). We found that YvcI can hydrolyze both ppGpp and pppGpp. In addition to (p)ppGpp, we also tested the ability of YvcI to hydrolyze GTP and 8-oxo-GTP to sanitize guanosine nucleotide pool^26^. YvcI failed to hydrolyze either GTP or 8-oxo-GTP (data not shown). The inability of YvcI to hydrolyze GTP and the strong structural similarity between GTP and (p)ppGpp suggest that NahA is a specific (p)ppGpp hydrolase which requires its substrate to have pyrophosphate group on the 3’ end. However, unlike *B. subtilis* RelA, which hydrolyzes the 3’-pyrophosphate group from ppGpp and pppGpp to produce GDP and GTP, NahA hydrolyzed ppGpp and pppGpp to yield a single nucleotide species that migrated differently than GTP (**Figure S3a**) or GDP (**Figure S3b**). To determine the identity of YvcI’s (p)ppGpp hydrolysis product, we analyzed the sample by liquid chromatography coupled with mass spectrometry (LC-MS), and compared to a pGpp standard produced by *E. faecalis* SAS (RelQ) *in vitro* ^22^. The LC-MS profile revealed a peak of the same mass over charge ratio (m/z) as GTP but with a different retention time (11.75 min versus 11.15 min for GTP). The retention time is the same as the pGpp standard (**Figure 2b**), suggesting that the hydrolysis product of YvcI is pGpp.

The production of pGpp from pppGpp and ppGpp agrees with the NuDiX hydrolase function, inferring that pppGpp and ppGpp are hydrolyzed between the 5’-α and 5’-β phosphate groups to produce guanosine-5’-monophosphate-3’-diphosphate (pGpp). Therefore, we renamed the enzyme NahA (<u>N</u>uDiX <u>a</u>larmone <u>h</u>ydrolase A).

It is possible, although unlikely, that NahA hydrolyzes the 3’-phosphate to produce ppGp, which would run at the same retention time as pGpp in LC-MS. To distinguish these two possibilities, we analyzed the NahA cleavage products of [3’-β-^32^P]-pppGpp. If NahA cleaves the 5’-γ and β phosphates, the reaction would yield [3’-β-^32^P]-pGpp. In contrast, if NahA cleaves the 3’-β-phosphate and the 5’-γ-phosphate, the reaction would yield free ^32^P-phosphate. TLC analysis revealed that the radioactive ^32^P after NahA hydrolysis of [3’-β-^32^P]-pppGpp co-migrates with the pGpp nucleotide rather than the free phosphate that migrates to the very end of TLC plate (**Figure 2c**). This result showed that the product has an intact 3’-pyrophosphate group, confirming the product to be pGpp rather than ppGp. Finally, quantification of [3’-β-^32^P]-pppGpp hydrolysis by NahA showed that the decrease of substrate radioactivity mirrored the increase of the single product radioactivity (**Figure 2d**), demonstrating the product is exclusively pGpp.

As a NuDiX hydrolase, NahA has been reported to have a modest activity in removing the 5’-phosphate of mRNA ^27^. We found that NahA is far more efficient at hydrolyzing (p)ppGpp than at decapping mRNA. Enzymatic assays revealed that NahA hydrolyzes ppGpp following Michaelis-Menten kinetics, with a k_cat_ of 1.22 ± 0.17 s^−1^ and a *K_m_* of 7.5 ± 2.3 μM (**Figure 2e, Table 1**). NahA also effectively and cooperatively hydrolyzes pppGpp, with a Hill coefficient of 2.78 ± 0.47, *k_cat_* of 10.0 ± 0.5 s^−1^ and *K_m_* of 177.4 ± 0.4 μM (**Figure 2e, Table 1**). In contrast, its *k_cat_* to decap RNA is around 0.0003 s^−1^ (estimation based on published figure) ^27^. The difference in *k_cat_ in vitro* suggest that NahA’s major function is to regulate (p)ppGpp rather than to decap the 5’ cap of mRNA.

**Table 1.** Kinetic parameters of (p)ppGpp hydrolysis by NahA.

|  | ppGpp | pppGpp |
| --- | --- | --- |
| $k_{cat}$ (s<sup>-1</sup>) | 1.22 ± 0.17 | 10.0 ± 0.5 |
| $K_M$ (μM) | 7.5 ± 2.3 | 177.4 ± 0.4 |
| Hill coefficient | 1 | 2.78 ± 0.47 |

### NahA hydrolyzes (p)ppGpp to produce pGpp *in vivo*

NahA was previously identified as a constitutively expressed protein with ~600 copies per cell ^28^. To examine its impact on (p)ppGpp *in vivo*, we engineered a *nahA* deletion strain, and developed an LC-MS metabolome quantification for pppGpp, ppGpp and pGpp in *B. subtilis* cells (see Material & Methods). LC-MS measurement of cell extracts showed that Δ*nahA* mutant cells accumulate more pppGpp and ppGpp than wild type cells during both log phase and stationary phase, in agreement with NahA’s ability to hydrolyze (p)ppGpp (**Figure 3a**). In contrast, Δ*nahA* mutant has much less pGpp than wild type cells (**Figure 3a**). Specifically, during log phase, pGpp can hardly be detected in Δ*nahA* (**Figure S4a**). When we complement Δ*nahA* with an overexpressed copy of *nahA*, the pGpp levels increased to more than wild type levels (**Figure S4**). These results support the function of NahA in producing pGpp.

**Figure 3.**
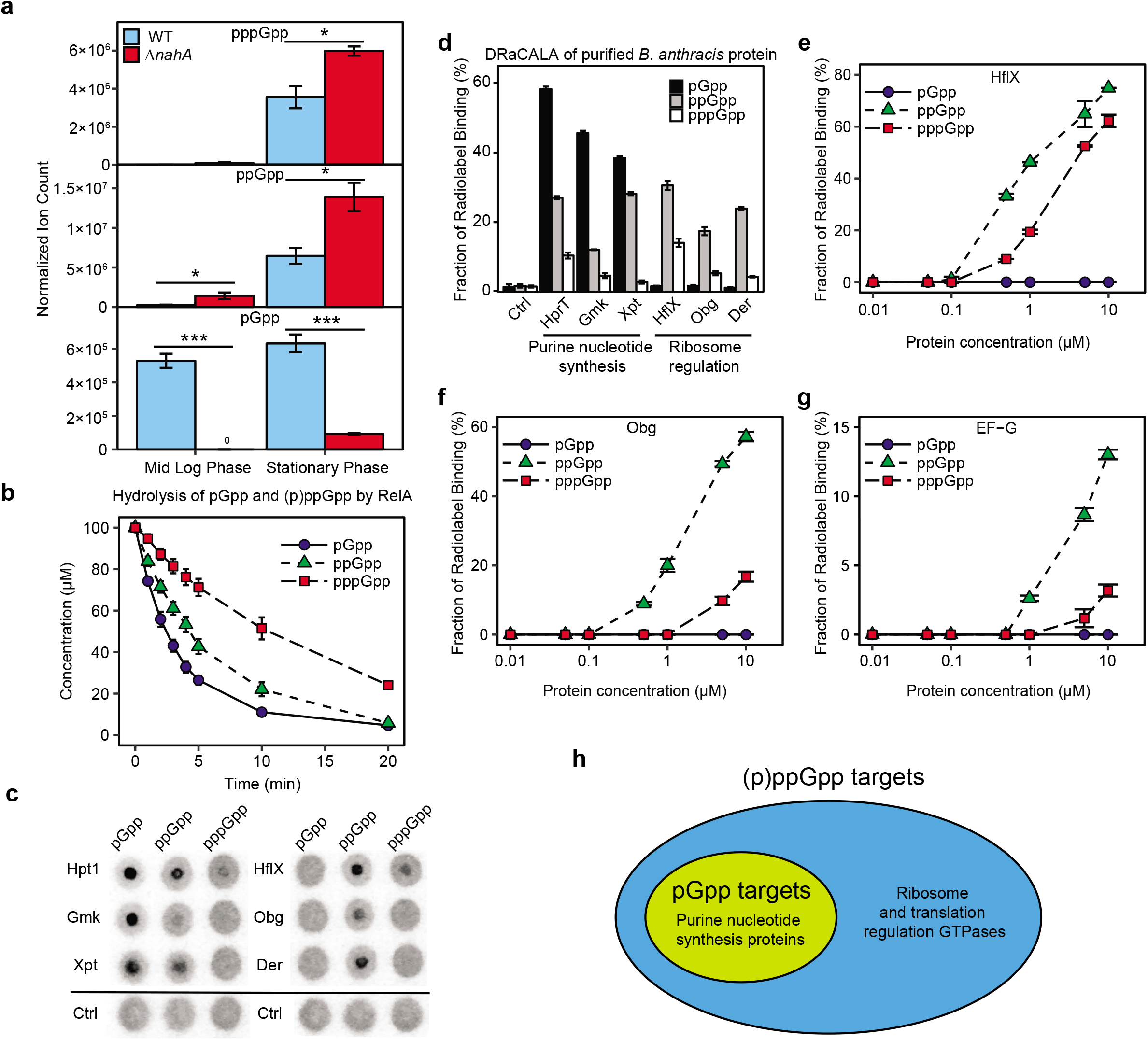
pGpp is produced by NahA *in vivo* and has a distinct binding spectrum compared to (p)ppGpp. **a**) LC-MS analyses of pGpp, ppGpp and pppGpp of wild type and Δ*nahA* in log phase and stationary phase. Normalized ion count is ion count per OD_600_ per unit volume of the culture. Error bars represent standard error of the mean of three biological replicates. A two-tailed two-sample equal-variance Student’s *t* test was performed between samples indicated by asterisks. Asterisks indicate statistical significance of differences (* *P*≤ 0.05, ** *P*≤ 0.01, *** *P*≤ 0.001). **b**) RelA hydrolysis of pGpp, ppGpp and pppGpp over time. For each nucleotide, 200 nM RelA was incubated with a mix of 100 μM unlabeled alarmone and a small amount of 5’-α-^32^P-labeled version. The degradation of the alarmone was analyzed by TLC to measure the decreased radioactivity. Error bars represent standard error of the mean of three replicates. **c**) DRaCALA of purified His-MBP-tagged *B. anthracis* proteins (1 μM) with less than 0.1 nM of identical amount of 5’-α-^32^P-labeled pGpp, ppGpp, and pppGpp. **d**) Quantification of DRaCALA in (c). **e-g**) Titrations of *B. anthracis* GTPases and quantification of their binding to pGpp, ppGpp and pppGpp using DRaCALA. GTPases HflX **e**), Obg **f**) and translation elongation factor G (EF-G) **g**). Error bars represent standard errors of the mean. **h**) Schematic showing the relationship between pGpp binding targets and (p)ppGpp binding targets.

We also used the drug arginine hydroxamate (RHX) which mimics amino acid starvation to induce accumulation of (p)ppGpp^12^. Using both LC-MS and TLC, we observed rapid accumulation of (p)ppGpp after RHX treatment, with Δ*nahA* cells showing stronger (p)ppGpp accumulation than wild type cells (**Figure S5a-d**). We still observe the accumulation of pGpp in Δ*nahA* cells, although to a much less extent than wild type cells (**Figure S5e**), this remaining pGpp in Δ*nahA* cells is likely due to the function of the enzyme SAS1, which can synthesize pGpp from GMP and ATP ^22^.

### pGpp can be hydrolyzed efficiently by RelA *in vitro*

The strong reducing effect of NahA on (p)ppGpp levels suggests that (p)ppGpp are efficiently hydrolyzed to pGpp. However, pGpp must be eventually degraded. In *B. subtilis*, pppGpp and ppGpp are both hydrolyzed by RelA, which has a functional (p)ppGpp hydrolase domain. Therefore, we performed *in vitro* assays using purified *B. subtilis* RelA and pGpp, ppGpp or pppGpp. We found that all three nucleotides can be hydrolyzed efficiently by RelA, furthermore, pGpp was hydrolyzed more rapidly than (p)ppGpp (**Figure 3b**).

### Protein binding spectrum of pGpp is distinct from (p)ppGpp

To understand whether pGpp is just an intermediate of (p)ppGpp hydrolysis or is a *bona fide* alarmone with its own regulatory targets, we used DRaCALA to systematically screen the *B. anthracis* library for pGpp-binding targets (**Figure 1a-b, Table S1**). Our screen showed that pGpp binds strongly to multiple purine nucleotide synthesis enzymes (**Figure 1c**), but binds to none of the (p)ppGpp-binding ribosome and translation regulation GTPases (**Figure 1d**). We then purified selected pGpp and (p)ppGpp binding targets and tested with [5’-α-^32^P]-labeled pGpp, ppGpp and pppGpp using DRaCALA (**Figure 3c-d**). These results confirmed strong pGpp binding to guanosine nucleotide synthesis proteins (Hpt1, Gmk, Xpt). In contrast, GTPases involved in ribosome biogenesis (HflX, Obg, Der) bind ppGpp strongly, but do not bind pGpp (**Figure 3d**). Titration analysis of these GTPases showed their strong affinity to ppGpp, modest affinity to pppGpp and lack of affinity to pGpp (**Figure 3e-g, Table S2**). We conclude that among the two main groups of conserved interaction targets of (p)ppGpp, pGpp exclusively regulates the purine pathway, but not the GTPases, thus can serve as a specialized signal (**Figure 3h**).

### *nahA* mutant exhibits stronger inhibition of translation upon amino acid starvation, delayed outgrowth and loss of competitive fitness

Finally, we analyzed the effects of *nahA* on cellular growth and metabolism. We first compared the key metabolites (NTPs, NDPs, NMPs, nucleosides and nucleobases) between wild type cells and the *nahA* mutant, using LC-MS analyses of expontential growth and stationary phase cells (**Figure 4a, Table S3**). Despite much higher (p)ppGpp levels and much lower pGpp levels in the *nahA* mutant, most purine nucleotides exhibit very little difference, corroborating with the observation that pGpp regulates purine metabolism similarly to (p)ppGp.

**Figure 4.**
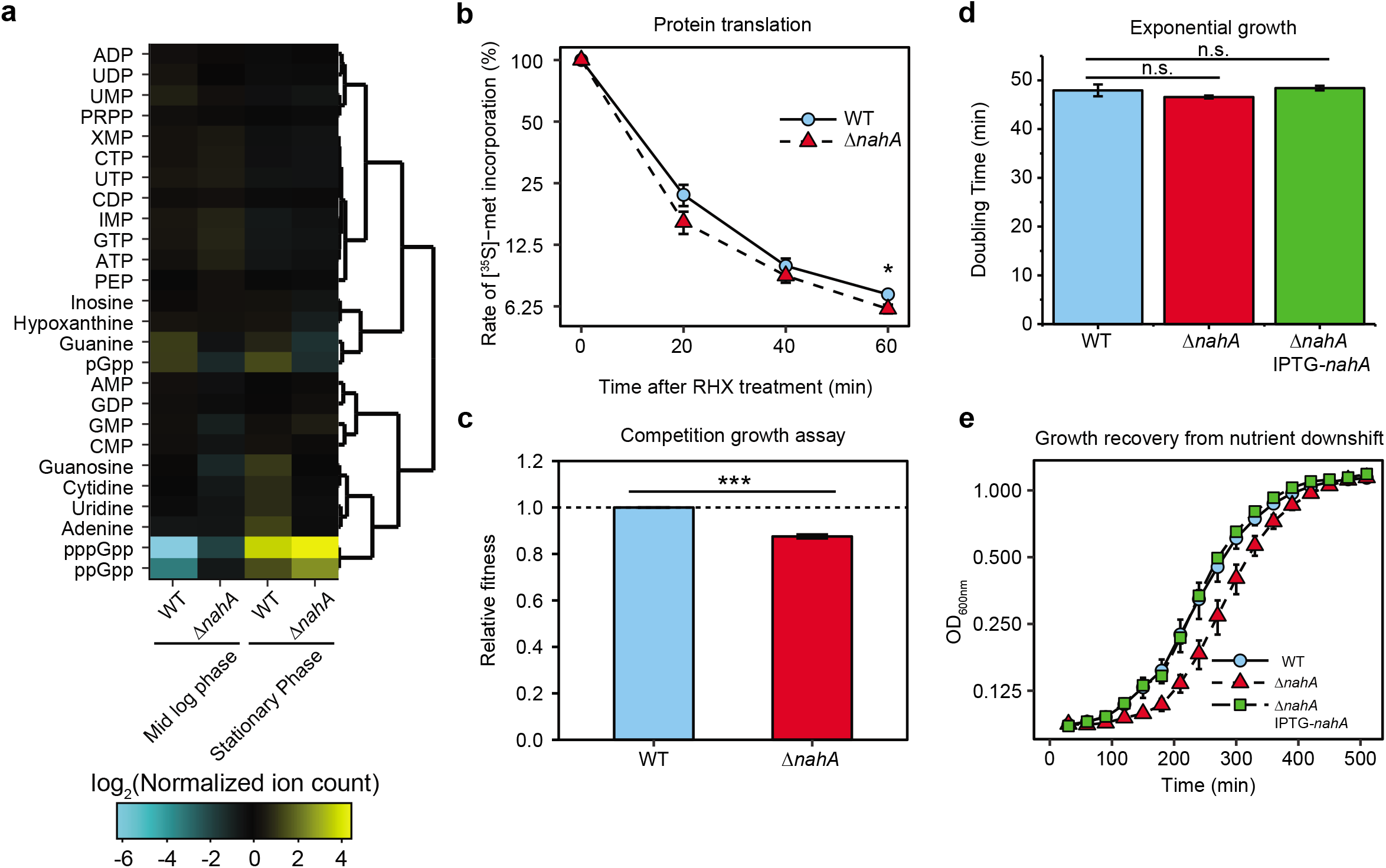
The effect of NahA on purine nucleotides, translation, fitness and growth recovery of *B. subtilis* cells. **a**) Hierarchical clustering of selected metabolites in exponential growth and stationary phase wild type and Δ*nahA* cells. Metabolites were measured by LC-MS. Normalized ion count is ion count per OD_600_ per unit volume of the culture. **b**) Protein translation rate of wild type and Δ*nahA* cells in a time course of arginine hydroxamate (RHX) treatment. Translation rate was measured by a two minutes pulse of [^35^S]-methionine incorporation into TCA-precipitable fraction. Error bars represent standard errors of the mean from three replicates. **c**) Relative fitness of Δ*nahA* and wild type strains obtained from a 7 day serial-dilution competition assay. Error bar represents standard deviation from 3 repeats. **d**) Doubling times of Δ*nahA* and wild type cells in log phase. A two-tailed two-sample equal-variance Student’s *t* test was performed between samples indicated by asterisks. Asterisk indicates statistical significance of difference (* *P*≤ 0.05, *** *P*≤ 0.001, n.s. *P*>0.05). **e**) Growth recovery from nutrient downshift. Log phase cultures of wild type, Δ*nahA* and *nahA* complementation (Δ*nahA* IPTG-*nahA*) strains in rich media were firstly downshifted in minimum media without amino acids for 10 min, and then diluted in rich media for outgrowth. Error bars in (d-e) represent standard errors from 6 replicates.

Next, we examined the effect of *nahA* on protein translation. The fact that pGpp does not directly regulate ribosome biogenesis and translation implicates an *in vivo* function of NahA: reducing (p)ppGpp levels to alleviate translation inhibition upon stress, while still keeping purine biosynthesis in check. To test this hypothesis, we measured total protein translation rate of wild type and Δ*nahA B. subtilis* by pulsed incorporation of ^35^S-methionine (**Figure 4b**). We found that after amino acid starvation, the rate of ^35^S-methionine incorporation is significantly further reduced in Δ*nahA* than in wild type (P<0.05), indicating a stronger inhibition of translation (**Figure 4b**).

Finally, we examined the impact of *nahA* on growth fitness of *B. subtilis*. We performed a growth competition assay in which a mixture of Δ*nahA* and wild type cells were grown to saturation and then diluted repeatedly for days. The proportion of Δ*nahA* rapidly decreased, indicating significant loss of fitness. The calculated relative fitness of Δ*nahA* is 0.875 ± 0.008 (mean ± S.D.) compared to wild type cells (**Figure 4c**). We found that Δ*nahA* has a similar doubling time as wild type cells (**Figure 4d**), but has a longer lag phase in adjusting to starvation (**Figure 4e**), likely due to excessive inhibition of translation (**Figure 4b**). Together, these results suggest that NahA tunes alarmone composition and alarmone levels to promote *B. subtilis* adjustment to nutrient fluctuation and optimizes growth and fitness.

## Discussion

Alarmones are universal stress signaling nucleotides in bacteria, however, the repertoire of alarmones and the target spectrums for each alarmone are incompletely understood. In this work, we have comprehensively characterized the interactomes of the related alarmones pppGpp and ppGpp and established the *in vivo* presence of pGpp as a closely related alarmone in Gram-positive *Bacillus* species. We characterized the direct targets of (p)ppGpp by screening an open reading frame expression library from *B. anthracis*. From this screen, we identified an enzyme NahA that converts (p)ppGpp to pGpp as efficient means to produce pGpp and to reduce (p)ppGpp concentrations, thus regulating the composition of the alarmones. We demonstrated that pGpp binds a distinct subset of protein receptors of (p)ppGpp. We also identified a key role of NahA in nutrient adaptation, suggesting that regulating alarmone composition may serve as a separation-of-function strategy for optimal adaptation.

### Conservation of pppGpp and ppGpp regulation of purine biosynthesis and ribosome biogenesis pathways across different species of bacteria

(p)ppGpp regulates diverse cellular targets that differ between different bacteria. For example, (p)ppGpp directly binds to RNA polymerase in *E. coli* to control the transcription of ribosomal and tRNA operons yet the (p)ppGpp binding sites on RNA polymerase are not conserved beyond proteobacteria^4–8^. Instead, (p)ppGpp accumulation in firmicutes strongly down-regulates synthesis of GTP, the exclusive transcription initiating nucleotides of rRNA and tRNA operons in firmicutes, to achieve a similar transcription control with different direct targets^12,29,30^. Therefore, identifying whether certain aspects of (p)ppGpp regulation are conserved among bacterial species is important for understanding bacterial survival and adaptation.

Our DRaCALA screen with a *B. anthracis* library revealed many ppGpp and pppGpp binding targets in this pathogenic Gram-positive bacterium. Comparison to ppGpp-binding targets in *B. anthracis* to *S. aureus*^18^ and *E. coli*^15,20^ identified novel targets but more importantly, revealed a clear theme of conservation. Most notably, (p)ppGpp in all three species binds to multiple proteins in two key pathways: 1) purine nucleotide synthesis, 2) translation-related GTPases including ribosome biogenesis factors.

The regulation of purine nucleotide synthesis in firmicutes includes well characterized targets Gmk and HprT whose regulation by (p)ppGpp protects *Bacillus subtilis* against nutrient changes like amino acid starvation and purine fluctuation, preventing cells from accumulating toxic high levels of intracellular GTP^12,14,31^. These also include new-found likely targets such as enzymes GuaC and PurA and the *de novo* pathway transcription factor PurR. Intriguingly, in the evolutionarily distant *E. coli*, the ppGpp targets in the purine biosynthesis pathway are different. For example, Gmk is not regulated by (p)ppGpp in *E. coli*. On the other hand, (p)ppGpp directly targets the *E. coli de novo* enzyme PurF^15^, a target that is not conserved in firmicutes. Therefore, despite differences in precise targets, (p)ppGpp extensively regulates the purine biosynthesis pathway in evolutionarily diverse bacteria (**Figure 1c**), highlighting this critical physiological role of (p)ppGpp.

We also found that (p)ppGpp interacts with essential GTPases that are implicated in ribosome biogenesis and translation in *B. subtilis* and *B. anthracis* (**Figure 1d**). (p)ppGpp’s targets in GTPases from *E. coli* to firmicutes are conserved^16–19^: HflX, Obg, Era, are also (p)ppGpp-binding proteins in *E. coli* and *S. aureus*^15,18,20^; Der and TrmE were identified in *E. coli* although not in *S. aureus* as (p)ppGpp-binding proteins^15,20^; RbgA doesn’t exist in *E. coli* and it is identified as a (p)ppGpp-binding protein in *S. aureus*^18^. HflX is a ribosome-spliting factor which may rescue stalled ribosomes under stressed conditions^32^, and mediates the dissociation of hibernating 100S ribosome to resume normal translation^33^. Obg^16^, Der^34^ and RbgA^35,36^ participate in the maturation of the 50S subunit of ribosome. Era functions in the assembly of the functional 70S ribosome complex^37^. TrmE functions in the maturation of tRNA to facilitate translation^38^. (p)ppGpp can be produced by amino acid starvation-induced translational stress or by defects in tRNA maturation^39^. Thus the conservation of (p)ppGpp regulation of GTPase targets highlights the key function of (p)ppGpp in quality control of ribosome biogenesis and regulation of translation.

### The third major alarmone pGpp in Gram positive species and its specific regulatory effect

In addition to pppGpp and ppGpp, it was long suspected that in *B. subtilis*, there are additional alarmones such as pppApp, pGpp, and ppGp accumulating during the stringent response^40,41^. Here we detected pGpp in *B. subtilis* cells, characterized an enzyme for its production *in vivo* and *in vitro*, and identified its targets. In contrast to our DRaCALA screens for pppGpp and ppGpp which identify similar targets and functionalities between these two alarmones, DRaCALA screen for pGpp displays a strong difference from (p)ppGpp with regarding to two conserved pathways. The affinity of pGpp for purine nucleotide synthesis proteins is equivalent or higher than that of (p)ppGpp. In contrast, pGpp’s affinity to GTPases involved in translational regulation is much lower, or absent, compared to (p)ppGpp. The distinct profile of target receptors distinguish pGpp as a different alarmone from (p)ppGpp, potentially allowing fine tuning of bacterial stress response. We propose a model for the function of NahA in growth recovery and competitive fitness by its role in transforming the alarmones. In wild type cells, (p)ppGpp produced in response to amino acid starvation will be hydrolyzed in part by NahA to pGpp. (p)ppGpp concentration in wild type cells during amino acid starvation can reach >1 mM. Given the copy number of NahA as ~600 copies per cell and its maximum velocities for (p)ppGpp hydrolysis, (p)ppGpp concentrations observed in wild type cells is a balance of RelA synthesis and NahA hydrolysis. Thus, in Δ*nahA* cells, (p)ppGpp accumulates higher than in wild type cells during amino acid starvation. Stronger inhibition of translation due to overly high level of (p)ppGpp in Δ*nahA* leads to slower growth recovery after culture enduring nutrient downshift and thus leads to fitness loss when co-cultured with wild type strain. In addition, because pGpp appears to be hydrolyzed more efficiently by RelA (**Figure 3b**), another role of NahA is perhaps to speed up (p)ppGpp removal to promote growth recovery.

In *E. coli*, (p)ppGpp binding proteins also include NuDiX hydrolases NudG and MutT^20^. Like NahA, NudG and MutT also hydrolyze (p)ppGpp. Unlike NahA, NudG and MutT produce guanosine 5’-monophosphate 3’-monophosphate (pGp) rather than pGpp. Similarly, in the bacterium *Thermus thermophilus*, the NuDiX hydrolase Ndx8 is also found to hydrolyze (p)ppGpp to produce pGp^42^. Sequence alignment shows that their homology with NahA is mostly restricted in the NuDiX box that is shared by NuDiX hydrolase family (**Figure S1**). Therefore, NudG, MutT and Ndx8 are considered alternative (p)ppGpp hydrolysis pathways to remove (p)ppGpp and promote growth^42^, rather than an alarmone-producer.

In *E. coli*, the enzyme GppA converts pppGpp to ppGpp, which may also regulate the composition of its alarmones^21^. In the absence of GppA, the alarmone pppGpp accumulates to a higher level than ppGpp^43^. It is possible that a similar separation-of-function regulation exists in *E. coli* by tuning pppGpp vs ppGpp levels. This can be addressed by examining pppGpp interactome in *E. coli*, which may reveal a differential theme than ppGpp. Ultimately, discovery of more enzymes that interconvert signaling molecules may reveal a universal theme of optimization through fine-tuning in signal transduction.

## Acknowledgement

We thank Mona Orr for technical help. We thank José Lemos for the gift of pGpp standard. This work is supported, in part, by an R35 GM127088 from NIGMS (to JDW), a GRFP DGE-1256259 from NSF (to BWA) and R01AI110740 from NIAID and NIDDK (to VTL).

## Materials and Methods

### *Bacillus anthracis* ORFeome Library Construction

*Bacillus anthracis* Gateway^®^ Clone Set containing plasmids bearing *B. anthracis* open reading frames was acquired from BEI Resources and used for Gateway cloning (Invitrogen protocol) into overexpression vectors pVL791 (10xHis tag ampicillin-resistant) and pVL847 (10xHis-MBP tag, gentamycin-resistant) and transformed into *Escherichia coli* BL21 lacI^q^ to produce two open reading frame proteome over-expression libraries (ORFeome library). The resulting ORFeome library contains 5139 ORFs from the genome of *B. anthracis str. Ames* (91.2% of 5632 ORFs in the genome). The corresponding proteins were expressed in *E. coli* and the cells were lysed to prepare the overexpression lysates for the downstream analysis.

### Plasmid and Strain Construction and Growth Conditions

Plasmid for NahA purification was constructed as follows: *nahA* encoding sequence was PCR amplified and incorporated into pE-SUMO vector by Golden Gate assembly method (New England BioLabs), and into pVL847 (His-MBP-tag overexpression vector) by Gateway cloning method (Invitrogen). Δ*nahA* mutant of *Bacillus subtilis* was constructed by CRISPR/Cas9 editing method (Ref). Primers for plasmid construction and mutation verification are listed in (**Table S4**). *nahA* deletion was confirmed by DNA sequencing.

If not specifically mentioned, *B. subtilis* strains were grown in S7 defined media ^44^ with modifications (50 mM MOPS instead of 100 mM, 0.1% potassium glutamate, 1% glucose, no additional tryptophan or methionine), with shaking at 250 rpm 37°C. The modified S7 defined media with 20 amino acids contains 50 μg/mL alanine, 50 μg/mL arginine, 50 μg/mL asparagine, 50 μg/mL glutamine, 50 μg/mL histidine, 50 μg/mL lysine, 50 μg/mL proline, 50 μg/mL serine, 50 μg/mL threonine, 50 μg/mL glycine, 50 μg/mL isoleucine, 50 μg/mL leucine, 50 μg/mL methionine, 50 μg/mL valine, 50 μg/mL phenylalanine, 500 μg/mL aspartic acid, 500 μg/mL glutamic acid, 20 μg/mL tryptophan, 20 μg/mL tyrosine, and 40 μg/mL cysteine.

### Nucleotide Preparation

(p)ppGpp was synthesized *in vitro* using Rel*Seq*_1-385_ and GppA, purified and quantified as demonstrated^21^. pGpp was synthesized, purified and quantified as demonstrated^22^.

Radiolabeled (p)ppGpp was synthesized *in vitro* using Rel*Seq*_1-385_ and GppA as demonstrated^21^ with modifications. We replaced non-radiolabeled GTP with 750 μCi/mL ^32^P-α-GTP (3000 mCi / mmol; PerkinElmer) for production of [5’-α-^32^P]-(p)ppGpp. We replaced non-radiolabeled ATP with 750 μCi/mL ^32^P-γ-ATP (3000 mCi/mmol; PerkinElmer) for production of [3’-β-^32^P]-ppGpp. After the reaction was done, we 1:10 diluted the reaction mix in Buffer A (0.1 mM LiCl, 0.5 mM EDTA, 25 mM Tris pH 7.5). The mixture was loaded onto a Buffer-A-equilibrated 1 mL HiTrap QFF column (GE Healthcare). Then the column was washed sequentially by 10 column volumes of Buffer A, and 10 column volumes of Buffer A with 170 mM LiCl (for ppGpp purification, the LiCl concentration was 160 mM) with 1 mL fractions. Radiolabeled (p)ppGpp was eluted by 5 column volumes of Buffer A with 500 mM LiCl with 1 mL fractions. The radioactivity and purity of radiolabeled (p)ppGpp were analyzed by thin-layer chromatography and phosphorimaging.

Radiolabeled pGpp was synthesized *in vitro* as demonstrated above for radiolabeled ppGpp synthesis with modifications. We replaced non-radiolabeled GTP with 750 μCi/mL ^32^P-α-GTP (3000 mCi / mmol; PerkinElmer) for production of [5’-α-^32^P]-pGpp. We replaced GppA with NahA for production of [5’-α-^32^P]-pGpp. After the reaction was done, we 1:10 diluted the reaction mix in Buffer A (0.1 mM LiCl, 0.5 mM EDTA, 25 mM Tris pH 7.5). The mixture was loaded onto a Buffer-A-equilibrated 1 mL HiTrap QFF column (GE Healthcare). Then the column was washed by 10 column volumes of Buffer A, and eluted by 10 column volumes of Buffer A with 155 mM LiCl with 1 mL fractions. Purified radiolabeled pGpp often come out in the 5^th^ to 9^th^ column volumes during elution. The radioactivity and purity of radiolabeled pGpp were analyzed by thin-layer chromatography and phosphorimaging.

### Overexpression and Purification of NahA

pE-SUMO-NahA was transformed into E. coli BL21(DE3) lacI^q^ by chemical transformation. A single colony of the corresponding strain was grown in LB with 30 μg/mL kanamycin overnight. Overnight culture was 1:100 diluted in LB-M9 media (7 g/L Na_2_HPO_4_, 2 g/L KH_2_PO_4_, 0.5 g/L NaCl, 1 g/L NH4Cl, 2 g/L glucose, 1 g/L sodium succinate dibasic hexahydrate; 10 g/L tryptone, 5 g/L yeast extract and 3 mM MgSO_4_) with 30 μg/mL kanamycin and grown at 30°C with shaking for 4 hr. After adding 1 mM IPTG for induction, the culture was further grown at 30°C for 4 hr. Then the culture was centrifuged at 4000 g for 30 min at 4°C and the supernatant was discarded. The pellet was stored at −80°C before cell lysis. All the following steps were performed at 4°C. Pellet was re-suspended in Lysis Buffer (50mM Tris-HCl pH 8, 10% sucrose w/v, 300mM NaCl) and lysed by French press. Cell lysate was centrifuged at 15000 rpm for 30 min, and the supernatant was collected, filtered through 0.45 μm pore-size cellulose syringe filter. His-SUMO-NahA filtered supernatant was purified using a 5 mL HisTrap FF column (GE Healthcare) equipped on an ÄKTA FPLC apparatus (GE Healthcare). SUMO Buffer A (50 mM Tris-HCl, 25 mM imidazole, 500 mM NaCl, 5% glycerol v/v) was used as washing buffer, and SUMO Buffer C (50 mM Tris-HCl, 500 mM imidazole, 500 mM NaCl, 5% glycerol v/v) was used as the elution buffer. Fractions containing most abundant His-SUMO-NahA were combined with 100 μg SUMO Protease and dialyzed twice against SUMO Protease Buffer (50 mM Tris-HCl, 500 mM NaCl, 5% glycerol v/v, 1 mM β-mercaptoethanol) overnight. His-SUMO tag was removed by flowing through HisTrap FF column and collecting the flow-through. Purified NahA was analyzed by SDS-PAGE and the concentration was measured by the Bradford assay (Bio-rad).

### Differential Radial Capillary Action Ligand Assay (DRaCALA)

Cell lysate with overexpressed protein and purified protein were used for DRaCALA as described^24^. 10 μL cell lysate or diluted, purified protein was mixed with 10 μL diluted [5’-α-^32^P]-(p)ppGpp or [5’-α-^32^P]-pGpp (~0.2 nM) in a buffer containing 10 mM Tris pH 7.5, 100 mM NaCl and 5 mM MgCl_2_, incubated at room temperature for 10 min. ~2 μL mixture was blotted onto nitrocellulose membrane (Amersham; GE Healthcare) and allowed for diffusion and drying. The nitrocellulose membrane loaded with mixture was exposed on phosphor screen, which was scanned by a Typhoon FLA9000 scanner (GE Healthcare). Fraction of (p)ppGpp or pGpp binding was analyzed as described^24^.

### Quantification of Intracellular Nucleotides by Thin-Layer Chromatography

Quantification of ATP, GTP, and (p)ppGpp levels by thin-layer chromatography was performed as described before ^45,46^ with modifications. Cells were grown in low phosphate S7 defined media (0.5 mM phosphate instead of 5 mM), labeled with 50 μCi/mL culture decontaminated ^32^P-orthophosphate (9000 mCi/mmol; Perkin-Elmer) (decontamination was performed as demonstrated previously^47^) at an optical density at 600 nm (OD_600nm_) of ~0.01 and grown for two to three additional generations before sampling. Nucleotides were extracted by adding 100 μL of sample into 20 μL 2 N formic acid on ice for at least 20 min. Extracted samples were centrifuged at 15,000 g for 30 min to remove cell debris. Supernatant samples were spotted on PEI cellulose plates (EMD-Millipore) and developed in 1.5 M or 0.85 M potassium phosphate monobasic (KH_2_PO_4_) (pH 3.4). TLC plates were exposed on phosphor screens, which were scanned by a Typhoon FLA9000 scanner (GE Healthcare). Nucleotide levels were quantified by ImageJ (NIH). Nucleotide levels are normalized to ATP level at time zero.

### Kinetic Assay of (p)ppGpp Conversion by NahA

100 μL reaction mix was prepared with 40 mM Tris HCl pH 7.5, 100 mM NaCl, 10 mM MgCl_2_, non-radioactive and ^32^P-radiolabeled (p)ppGpp (concentration as required by specific experiments), 100 nM purified NahA (add enzyme last). Before adding enzyme, reaction mix was prewarmed at 37°C for at least 10 min. After adding enzyme, 10 μL reaction mix was aliquoted into 10 μL ice-chilled 0.8 M formic acid to stop the reaction and preserve nucleotides at every time point. Samples at each time point were resolved by thin-layer chromatography on PEI-cellulose plates with 1.5 M KH_2_PO_4_ (pH 3.4). Nucleotide levels were quantified as mentioned above and the phosphorimager counts of substrate and product were used to calculate the concentration of product by a formula:

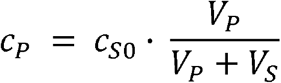

Here *c_P_* was the concentration of product, *c*_*S*0_ was the concentration of substrate before the reaction starts, *V_P_* was the phosphorimager count of product, and *V_S_* was the phosphorimager count of substrate. Initial rates of hydrolysis (*v*_0_) were calculated using slope of the initial linear part of *c_P_* over time curve at different initial substrate oncentrations. Michaelis-Menten constant (*K_m_*) and catalytic rate constant (*k_cat_*) were obtained by fitting the data of *v*_0_-*c*_*S*0_ to the model

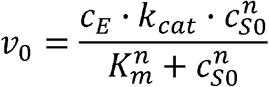

by MATLAB, where *c_E_* was the concentration of NahA, *c*_*S*0_ was the initial concentration of substrate, and *n* was the Hill’s coefficient (for ppGpp hydrolysis, fix *n* to 1).

### Kinetic Assay of (p)ppGpp Hydrolysis by RelA

100 μL reaction mix was prepared with 25 mM Tris HCl pH 7.5, 1 mM MnCl_2_, 100 μM non-radioactive and ~0.2nM ^32^P-radiolabeled (p)ppGpp, 200 nM purified RelA (add enzyme last). Before adding enzyme, reaction mix was prewarmed at 37 “C for at least 10 min. After adding enzyme, 10 μL reaction mix was aliquoted into 10 μL ice-chilled 0.8 M formic acid to stop the reaction and preserve nucleotides at every time point. Samples at each time point were resolved by thin-layer chromatography on PEI-cellulose plates with 1.5 M KH_2_PO_4_ (pH 3.4). Nucleotide levels were quantified as mentioned above and the phosphorimager counts of substrate and product were used to calculate the concentration of product by a formula:

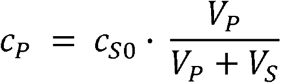

Here *c_P_* was the concentration of product, *c*_*S*0_ was the concentration of substrate before the reaction starts, *V_P_* was the phosphorimager count of product, and *V_S_* was the phosphorimager count of substrate.

### LC-MS Quantification of Metabolites

Cells were grown to designated OD_600nm_. 5 mL of cultures were sampled and filtered through PTFE membrane (Sartorius) at time points before and after 0.5 mg/mL arginine hydroxamate treatment. Membranes with cell pellet were submerged in 3 mL extraction solvent mix (on ice 50:50 (v/v) chloroform/water) to quench metabolism, lyse the cells and extract metabolites. Mixture of cell extracts were centrifuged at 5000 g for 10 min to remove organic phase, then centrifuged at 20000 g for 10 min to remove cell debris. Samples were frozen at −80°C if not analyzed immediately. Samples were analyzed using an HPLC-tandem MS (HPLC-MS/MS) system consisting of a Vanquish UHPLC system linked to heated electrospray ionization (ESI, negative mode) to a hybrid quadrupole high resolution mass spectrometer (Q-Exactive orbitrap, Thermo Scientific) operated in full-scan selected ion monitoring (MS-SIM) mode to detect targeted metabolites based on their accurate masses. MS parameters were set to a resolution of 70,000, an automatic gain control (AGC) of 1e6, a maximum injection time of 40 ms, and a scan range of 90-1000 mz. LC was performed on an Aquity UPLC BEH C18 column (1.7 μm, 2.1 × 100 mm; Waters). Total run time was 30 min with a flow rate of 0.2 mL/min, using Solvent A (97:3 (v/v) water/methanol, 10 mM tributylamine pH~8.2-8.5 adjusted with approximately 9 mM acetic acid) and 100% acetonitrile as Solvent B. The gradient was as follows: 0 min, 5% B; 2.5 min, 5% B; 19 min, 100% B; 23.5 min 100% B; 24 min, 5% B; 30 min, 5% B. Raw output data from the MS was converted to mzXML format using inhouse-developed software, and quantification of metabolites were performed by using the Metabolomics Analysis and Visualization Engine (MAVEN) software suite^48,49^. Normalized ion count was defined and calculated as the ion count per OD_600_ per unit volume (5 mL) of the culture.

### Growth Competition Assay

*nahA::erm^R^* mutant and wild type cells were mixed in LB broth to an OD_600nm_ of 0.03 and grown at 37°C with vigorous shaking. After every 24 hour period, the stationary phase culture was back-diluted in fresh LB broth to an OD_600nm_ of 0.02, for a total period of 7 days. Each day, the culture was sampled, serially diluted and spread over LB agar and LB agar containing 0.5 μg/mL erythromycin/12.5 μg/mL lincomycin to obtain the CFU of total bacteria and erythromycin-resistant strain, respectively. Relative fitness was calculated by the formula^50^: 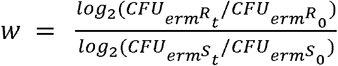, where *w* means relative fitness of the erythromycin-resistant strain to the erythromycin-sensitive strain; *CFU*_*erm^R^*_0__ and *CFU_erm^R^_t__* means the total CFU of erythromycin-sensitive strain before and after competition, respectively; *CFU*_*erm^S^*_0__ and *CFU_erm^S^_t__* means the total CFU of erythromycin-sensitive strain before and after competition, respectively.

### [^35^S]-Methionine Incorporation Assay

[^35^S]-Methionine (10.2 mCi/mL, PerkinElmer) was diluted to 0.5 μCi/μL and aliquoted into 1.5 mL Eppendorf tubes (10 μL per tube, heated at 37□). Cells were grown to OD_600nm_ at around 0.2 (record the exact OD). Aliquots of 200 μL culture were taken at time points before and after 0.5 mg/mL RHX treatment, added into tubes of aliquoted [^35^S]-Methionine, incubated at 37°C for 5 min and fixed by adding 200 μL ice-chilled 20% (w/v) trichloroacetic acid. Fixed samples were chilled before filtration. Glass fiber filters (24mm, GF6, Whatman) were prewetted with 3 mL ice-chilled 5%(w/v) trichloroacetic acid and applied with fixed samples. Filters were then washed three times by 3 x 10 mL ice-chilled 5%(w/v) trichloroacetic acid and dried by ethanol. Dried filters were put in scintillation vials, mixed with 5 mL scintillation fluid and then sent for scintillation count in the range of 2.0-18.6 eV. Counts per minutes (CPM) measured by scintillation counter were used as the representative of translation rate. The ratio of the CPM of one sample relative to the CPM at time zero was calculated as the relative translation rate.

### Growth recovery from nutrient downshift

Cells were grown in defined rich medium (S7 Glucose + 20 amino acids) to mid log phase, and then washed in poor medium (S7 Glucose without amino acid). Washed cultures were diluted into fresh rich medium and the growth was monitored by a plate reader (Synergy 2, Biotek) at 37°C under shaking.

---

**Figure S1.**
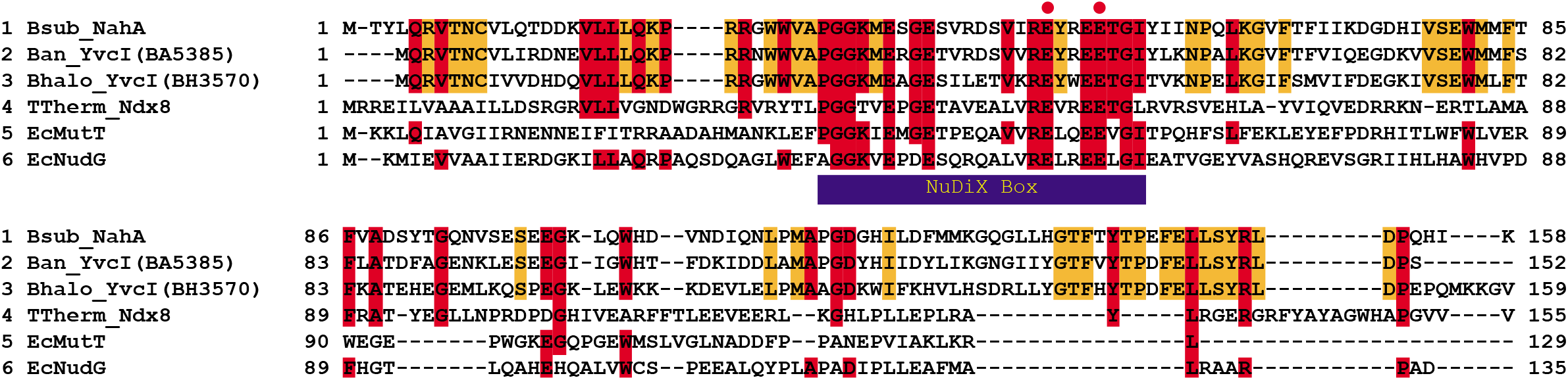
Protein sequence alignment of NahA (YvcI) homologs in different species. Multiple sequence alignment of YvcI homologs from *B. subtilis* (Bsub), *B. anthracis* (Ban), *B. halodurans* (Bhalo), *Thermus thermophilus* (TTherm) and *E. coli* (Ec). Alignment was obtained with MAFFT. Red shading indicates identical residues among *Bacillus* and other species, and orange shading indicates identical residues only in *Bacillus* species. The *Bacillus halodurans* YvcI structure (PDB ID: 3FK9) shows its conserved structure as a NuDiX hydrolase. Red dots label the key residues for catalytic activity. Numbers on the sides of the sequences indicate the residue numbers in the corresponding proteins.

**Figure S2.**
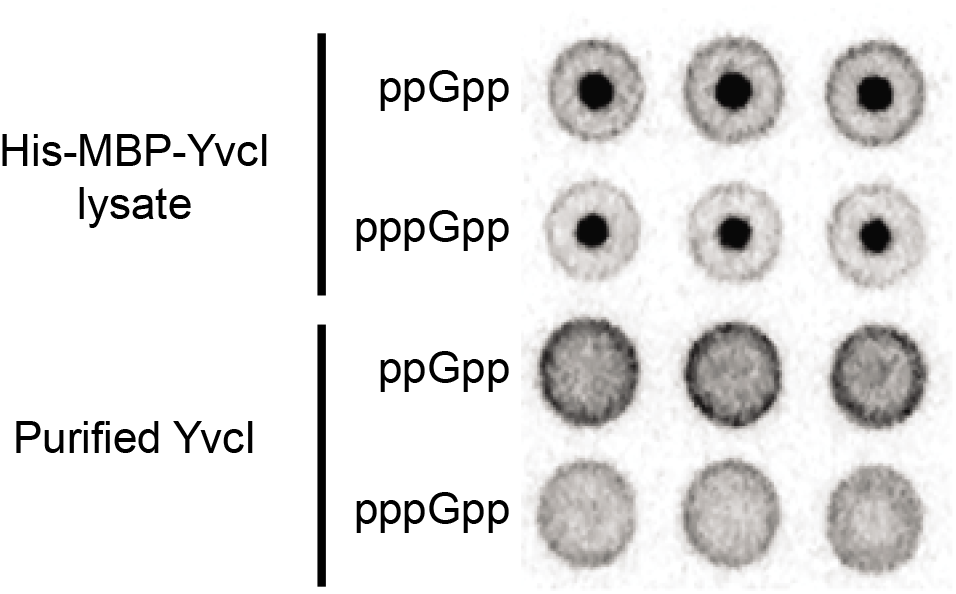
Cell lysate containing YvcI, but not purified YvcI, binds (p)ppGpp. DRaCALA of cell lysate containing His-MBP-tagged *B. subtilis* YvcI(NahA) and purified His-MBP-tagged *B. subtilis* YvcI(NahA) with 5’-α-^32^P-labeled (p)ppGpp.

**Figure S3.**
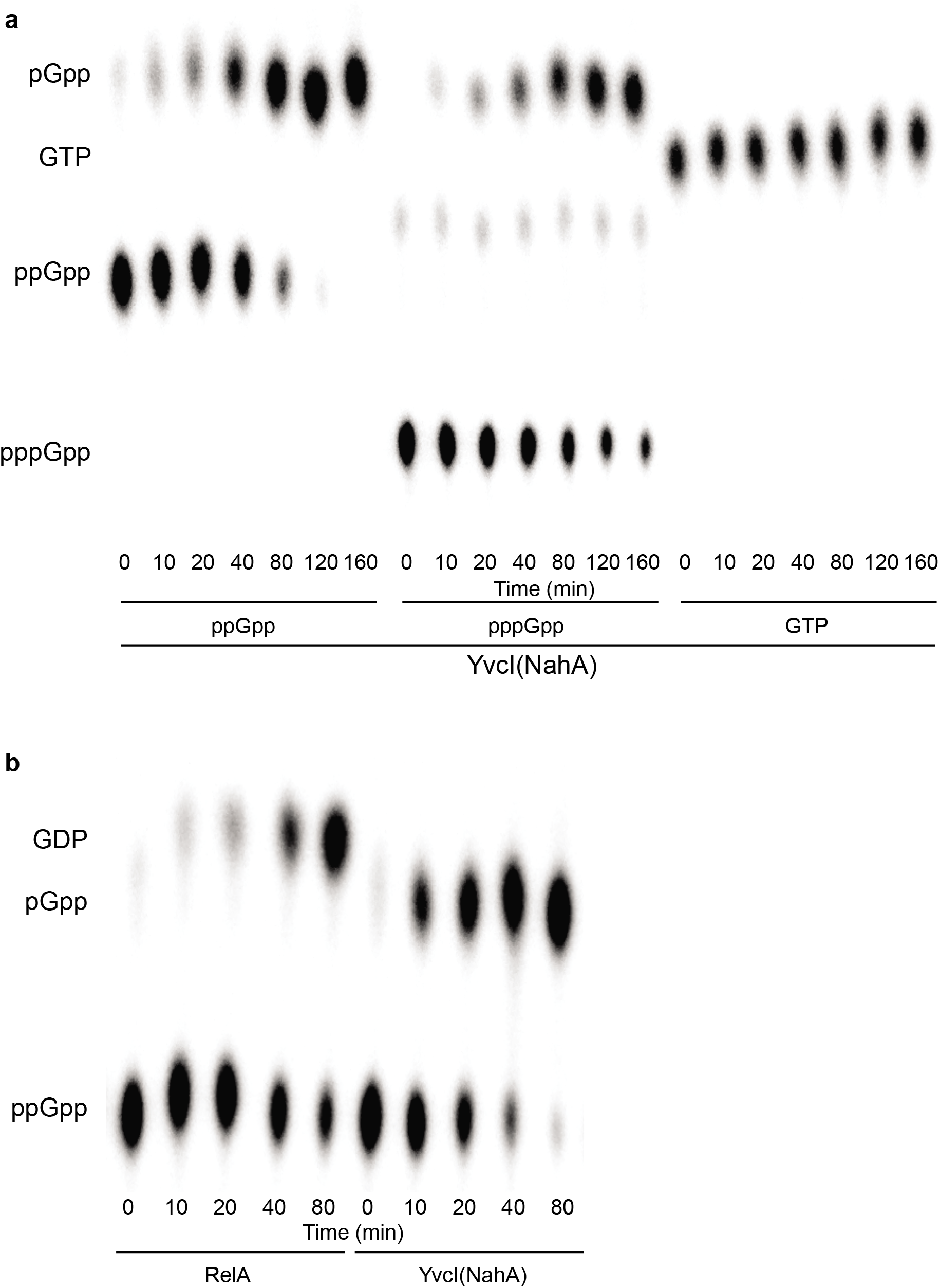
NahA hydrolyzes (p)ppGpp *in vitro*. **a**) TLC analysis of pppGpp and ppGpp hydrolysis by NahA over time. The same assay was also performed using GTP. **b**) TLC analysis of ppGpp hydrolysis by RelA and NahA over time. Substrate and product identities are labeled on the left side of the figure.

**Figure S4.**
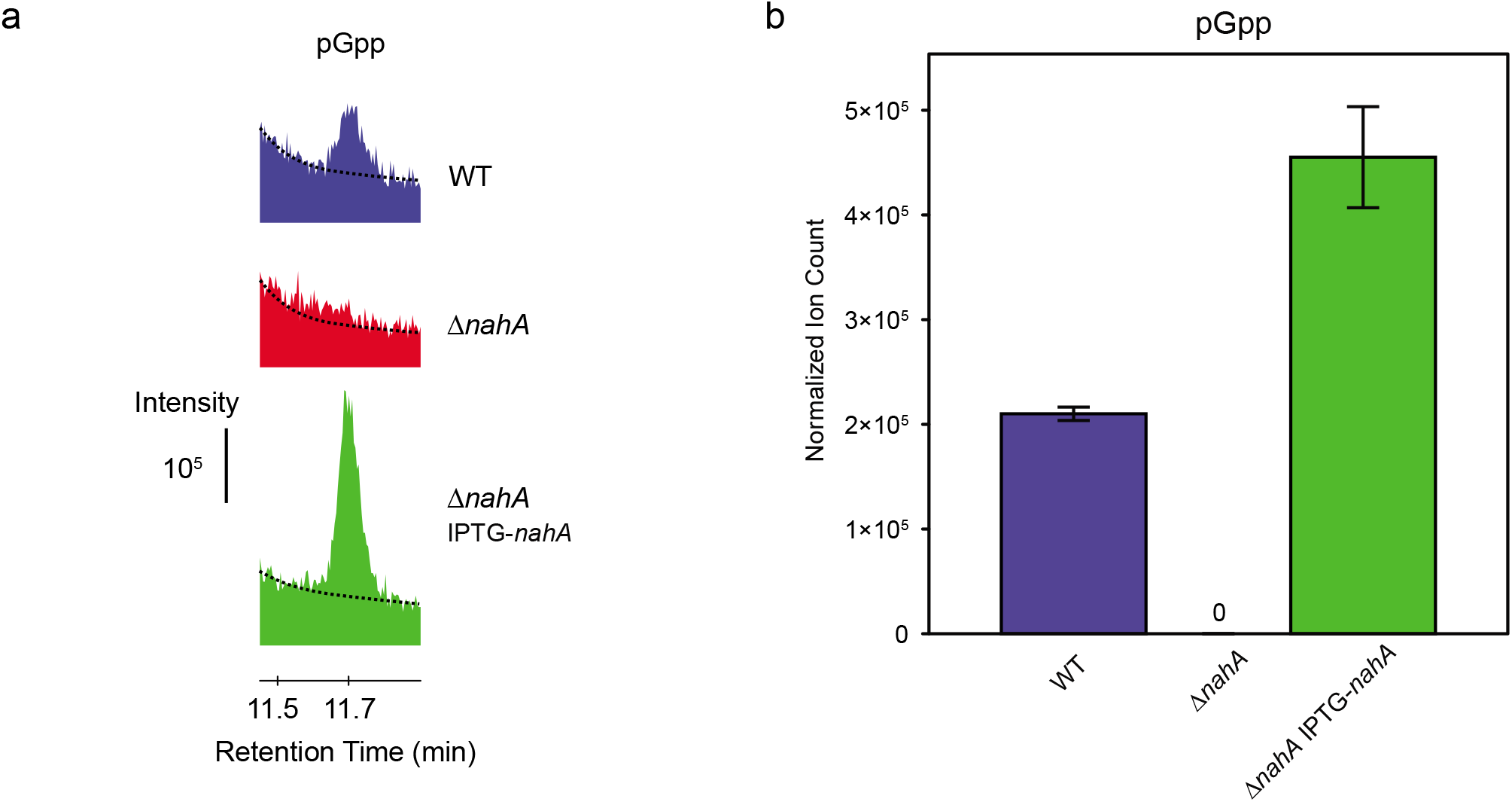
*nahA* complementation rescues pGpp production *in vivo*. **a**) Intensity profile of pGpp from LC-MS measurement of mid log phase wild type, Δ*nahA* and *nahA* complementation strain (Δ*nahA* IPTG-*nahA*). x-axis: retention time in LC-MS; y-axis: ion counts. For each strain, a representative curve was shown. Dashed lines are the shoulders of the GTP signal peak. **b**) Normalized ion count (ion count per OD_600_ per unit volume of the culture) of corresponding pGpp levels, obtained as the average of 3 biological replicates in **a**). Error bars represents standard error of the mean.

**Figure S5.**
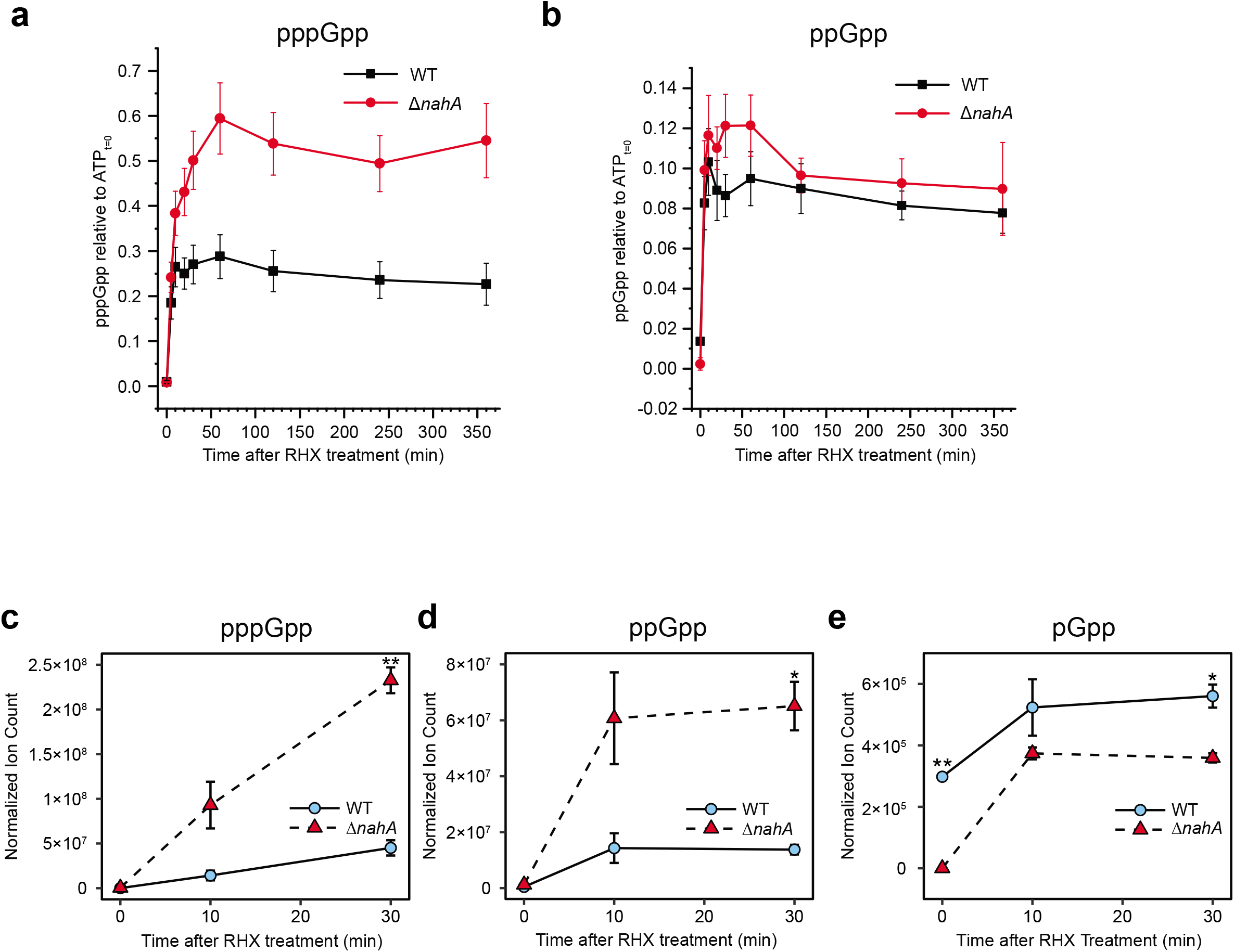
*nahA* mutant displayed higher (p)ppGpp and lower pGpp *in vivo* in *B. subtilis* in a time course of amino acid starvation. **a-b**) pppGpp (a) and ppGpp (b) levels in wild type and Δ*nahA B. subtilis* cells treated with the nonfunctional arginine analog RHX (arginine hydroxamate). Nucleotide levels were assayed via TLC and are normalized to the ATP level at time zero (ATP_t=0_). Error bars represent standard errors of mean of three independent experiments. **c-e**) LC-MS analyses of pppGpp (c), ppGpp (d) and pGpp (e) of wild type and Δ*nahA* in a time course of RHX treatment. Normalized ion count is ion count per OD_600_ per unit volume of the culture. Error bars represent standard error of the mean of two biological replicates. Asterisks indicate statistical significance of differences (* *P*≤ 0.05, ** *P*≤ 0.01).

