## Supplementary material for "Systemic characterization of pppGpp, ppGpp and pGpp targets in *Bacillus* reveals NahA converts (p)ppGpp to pGpp to regulate alarmone composition and signaling": Table S2

Table S2. *K_d_* ± SD of GTPases to pppGpp, ppGpp and pGpp

|  | pppGpp | ppGpp | pGpp |
| --- | --- | --- | --- |
| HflX | 5 ± 1 μM | 3.0 ± 0.6 μM | ND |
| Obg | 48 ± 7 μM | 6 ± 2 μM | ND |
| EF-G | 296 ± 50 μM | 60 ± 8 μM | ND |
